# Who is the host of the host-associated microbiome? Colony-level dynamics overshadow individual-level characteristics in the fur microbiome of a social mammal, the Egyptian fruit-bat

**DOI:** 10.1101/232934

**Authors:** Oren kolodny, Maya Weinberg, Leah Reshef, Lee Harten, Abraham Hefetz, Uri Gophna, Marcus W Feldman, Yossi Yovel

**Affiliations:** Department of Biology, Stanford University, Stanford; Department of Zoology, Tel Aviv University, Tel Aviv; School of Molecular Cell Biology and Biotechnology

**Keywords:** fruit bats, microbiome, pathogens, ecology, behavior

## Abstract

In the first longitudinal study of bat microbiomes, we find that unlike the pattern described in humans and other mammals, the prominent dynamics in Egyptian fruit bats’ fur microbiomes are those of change over time at the level of the colony as a whole. Thus, on average, a pair of fur microbiome samples from different individuals in the same colony collected on the same date are more similar to one another than a pair of samples from the same individual collected at different time points. This pattern suggests that the whole colony may be the appropriate biological unit for understanding some of the roles of the host microbiome in social bats’ ecology and evolution. This pattern of synchronized colony changes over time is also reflected in the profile of volatile compounds in the bats’ fur, but differs from the more individualized pattern found in the bats’ gut microbiome.

## Introduction

The host-associated microbiome may play a role in the host’s fate on many timescales, from the short-term health and behavior of the individual ((1–8)), through the life-long ecology and life history of the animal ((9)), to the long-term evolutionary adaptation of a species to its environment ((10–12)). However, data regarding non-human vertebrate microbiomes, particularly those of skin or fur, are just beginning to accumulate, and our understanding of the processes that determine their composition and function is limited. This is particularly true for non-model organisms: to date, few studies have collected longitudinal samples of non-human microbiomes in ecologically-realistic settings ((13)).

In this article we report on the temporal dynamics of the fur and the gut microbiome, assessed using 16S rRNA gene amplification, of ten Egyptian fruit bats (*Rousettus aegyptiacus*) in a captive colony (henceforth, the ‘experimental colony’). The colony was sampled weekly over a period of 13 weeks, in addition to sparser sampling of all 33 individuals in this colony and of a few additional individuals in a non-captive colony over the same time period (henceforth the ‘open colony’). We compare these data to the volatile components found in the bats’ fur, collected weekly and analyzed using gas chromatography (*GC*).

Our findings allow us to address an on-going debate regarding the conceptualization of host-microbiome ecology and evolution: some perspectives emphasize the potential utility of a holobiont theory, which regards the host and its associated microbial species together as a meaningful ecological and evolutionary unit (14, 15). Others focus on a metagenomic function-oriented account of the host and its associated microbiota (16, 17). Yet others suggest that no such novel theory is required, and that the ecology of the host and its associated microbiome can best be understood in terms of existing evolutionary and ecological theory, e.g. microbe-environment interaction or generalized multi-species Lotka-Volterra dynamics (18). This debate also recalls an earlier debate in evolutionary biology about levels of selection and ecological dynamics, with different perspectives suggesting the gene, the individual, the kin group, or the social group as the meaningful biological unit, whose trajectory in time is most useful and meaningful to track (19, 20). The different perspectives are not mutually exclusive, however, and may contribute complementary insights.

Our empirical findings in the bats’ fur microbiome suggest a perspective that to the best of our knowledge is new, and which informs both the conceptualization of the host-associated microbiome and the question of the level at which ecological dynamics take place. We propose that in some cases the whole colony of host organisms functions as a collective host with which the microbiome is associated.

A range of studies in humans and a few in wild animals have suggested that at certain body sites, such as the gut or skin, the primary determinant of the microbiome composition is individual identity (21–29). That is, on average, two microbiome samples from the same individual, taken at different time points, will be more similar to one another in their composition than two samples from different individuals, even for individuals controlled for sex, age, and other variables. Here we report that this regularity is not seen in the composition and dynamics of the fur microbiome of a highly social mammal that roosts in tight colonies – Egyptian fruit bats. Instead, we find that changes over time in the fur microbiome are best described as occurring at the colony level, with inter-individual variation playing a secondary role. The pattern seen in the gut microbiome, however, is different: some coordinated change in microbiome composition occurs, but this phenomenon is secondary to the role of individual identity and sex in determining individuals’ gut microbiomes.

Dynamics of change over time occur in the bats’ fur chemistry as well: the bats’ fur constitutes a habitat whose conditions strongly influence the composition of the microbiome, and are also affected by it. The idea that an animal’s microbiome will shape its odor and will thus play a role in its sociality (e.g., via olfactory recognition) has been raised multiple times, but studied mostly with respect to the microbiome in scent glands or specialized organs involved in olfactory communication ((30–36)). As with the composition of the microbiome, our results suggest a colony-level change over time of the bats’ fur chemistry. In addition, certain microbial taxa are linked to changes in the furs’ profile of volatile compounds.

## Results

A total of 518 samples of the fur and gut microbiota of bats, together with 36 samples of food and environmental control samples, were analyzed in this study, yielding 7196 non-chimeric Operational Taxonomic Units (OTUs) at 99% identity. These could be assigned (Figure 1) to 581 bacterial species, belonging mainly to the phyla Firmicutes (mean relative abundance in fur: 46%, and in gut: 39%) and Proteobacteria (mean relative abundance in fur: 30%, in gut: 38%). The near absence of Bacteroidetes, usually prevalent in animal guts (37), is noteworthy and likely related to the fruit bats’ unique habits and diet. To minimize weight during flight, bats chew the fruit, ingesting the juices and discharging much of the remaining pulp; ingested fruit passes rapidly through the digestive tract, with a short retention time of ~40 minutes (38, 39). The fruit bats’ diet thus relies heavily on consumption of simple sugars; accordingly, the intestine is relatively short, undifferentiated and with no observed cecum. It seems quite possible that this short time, and the non-compartmental structure of the digestive tract, do not allow appropriate conditions for the anaerobic fermentation of complex sugars that the Bacteroidetes typically carry out. Fur samples were far more diverse than gut samples (mean Shannon index 5.15, as opposed to 5.84, p<2.2e-16), yet there was also a high degree of overlap between the communities, with *Streptococcus salivarius* (mean abundance in fur: 16%, in gut: 13%) being the most common species in both sites. The microbiome composition in samples from the experimental colony was comparable to that of the open colony, a colony of fruit bats that roost in our cave-like facility, but behave as wild bats, flying nightly out to forage (Figure 1, and see below).

**Figure 1.**
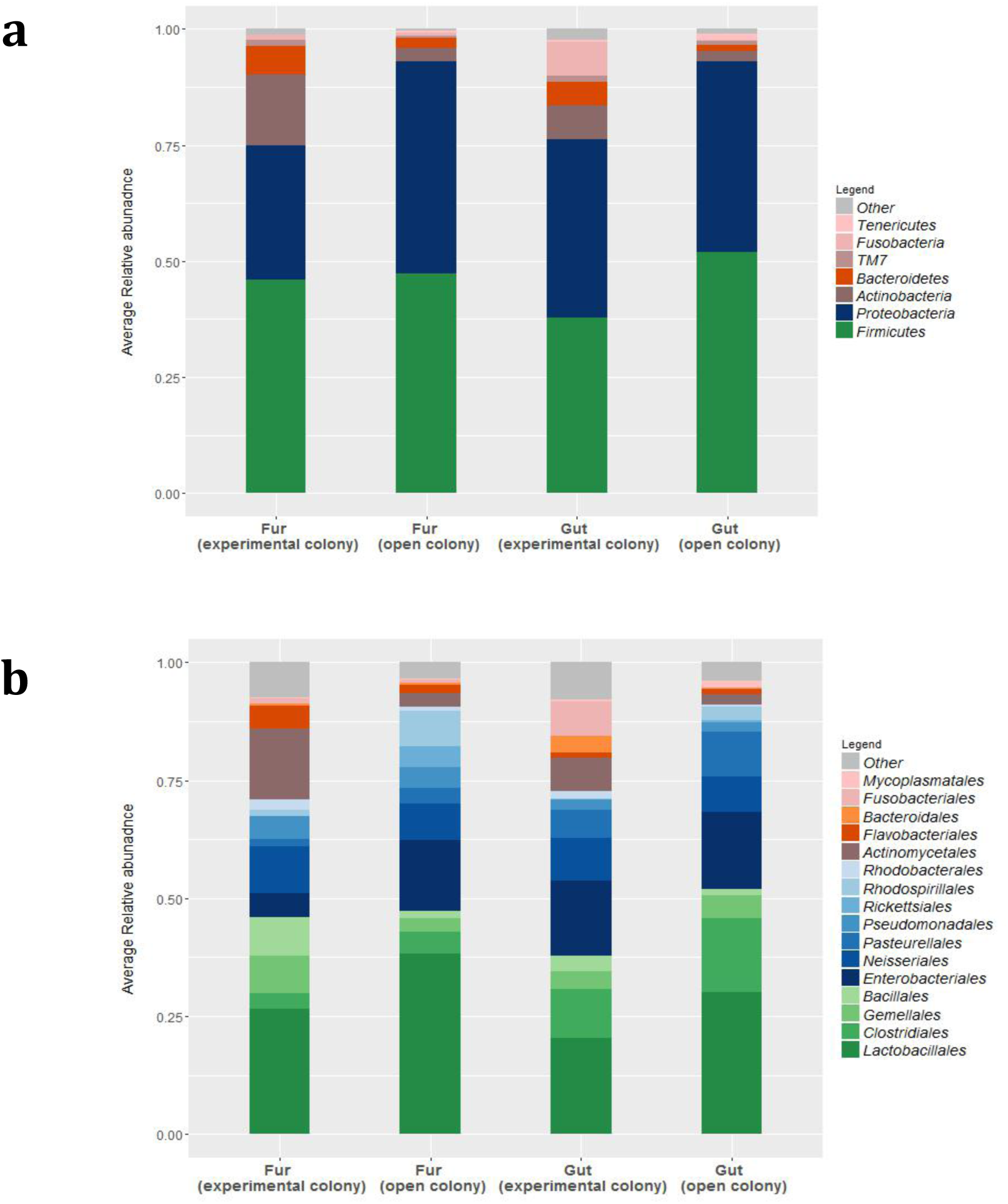
Microbial composition of fur and gut samples. The average relative abundance of each taxon per site is shown. All taxa with a mean relative abundance < 1% were grouped to a single category named “Other”. (a) Phylum level; (b) Order level; orders belonging to the Proteobacteria phylum are in shades of blue; orders belonging to the Firmicutes phylum are shown in shades of green.

The most striking observation, and the focus of this report, was a coordinated change in fur microbiome composition over time across all individuals (**Figure 2a**)^1^. This pattern is strong relative to the lack of visual clustering of the samples according to individual identity (**Figure 2b**). A permutation analysis of variance supports this visual observation: the date on which each sample was taken explains circa 35% of the fur microbiome variance in the experiment colony, while individual identity explains only circa 8% of the variance (PERMANOVA test using the Adonis method in R, p<0.001). To validate this finding, we conducted a comparison of the distances between pairs of samples. We found (**Figure 3a**) that samples from different individuals on the same date are, on average, more similar to one another than samples taken on different dates but from the same individual (Kruskal-Wallis test, p-value<0.001; confirmed using a Mantel test to avoid pseudo-replication, p-value<0.001; See analogous analyses with other distance measures in Supplementary Section 3).

**Figure 2:**
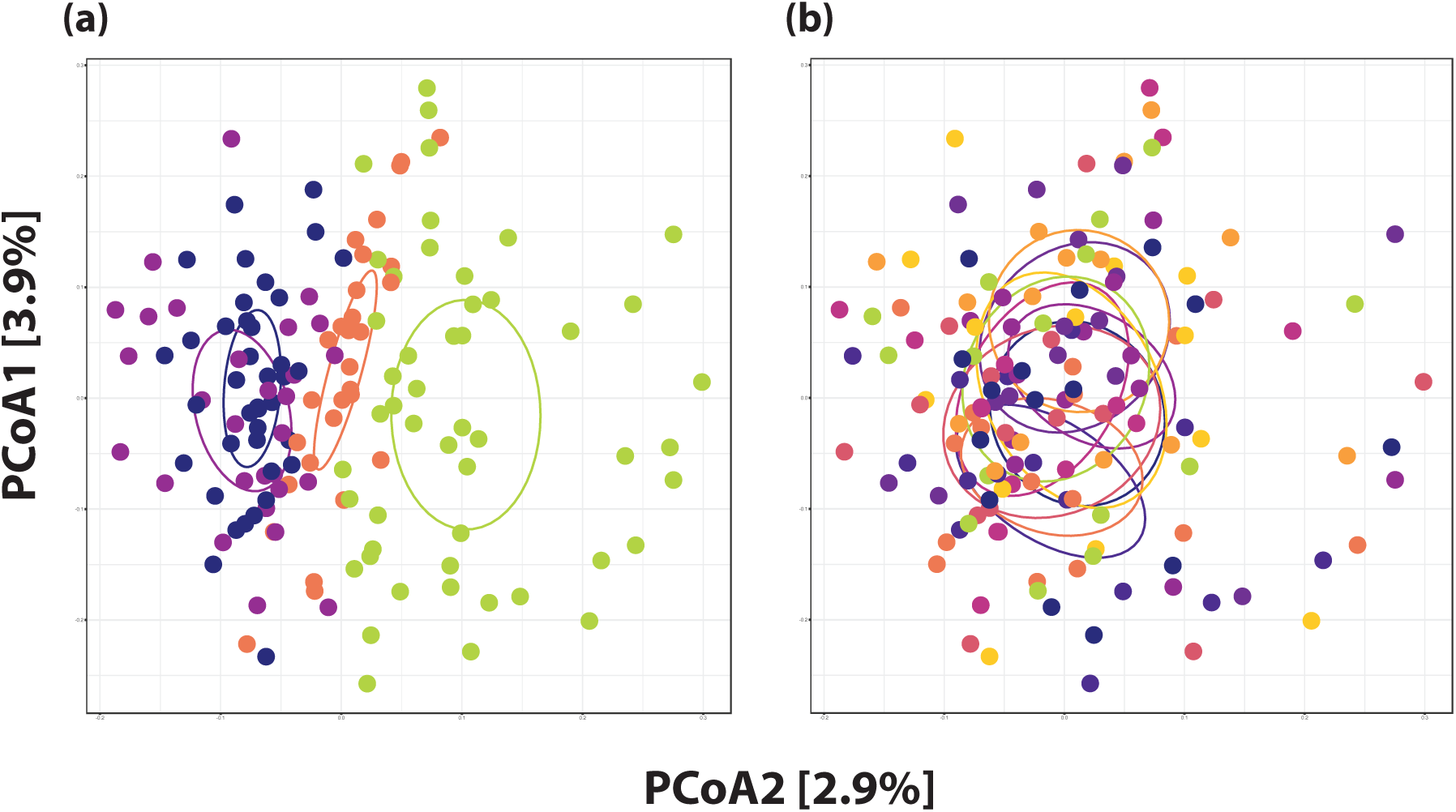
The prominent pattern in the fur microbiome is that of colony-level change over time. Samples from the fur of the ten focal bats in the experiment colony (weekly samples, over 13 weeks), plotted using PCoA of the Jaccard distance between samples. Each point represents a sample. In **(a),** each sample is colored according to its date of sampling; dates are divided into the four time quarters of the 13-week period of the experiment (quarters 1-4 are denoted by blue, purple, orange, and green, respectively). The clustering is seen to correspond to the quartile of sampling. In **(b)**, each sample is colored according to the individual bat from which it was taken. No clear clustering according to individual identity is visually apparent. Here and in all other PCoA plots, each ellipse represents the region around the center of mass of the samples in the group (see Methods).

**Figure 3:**
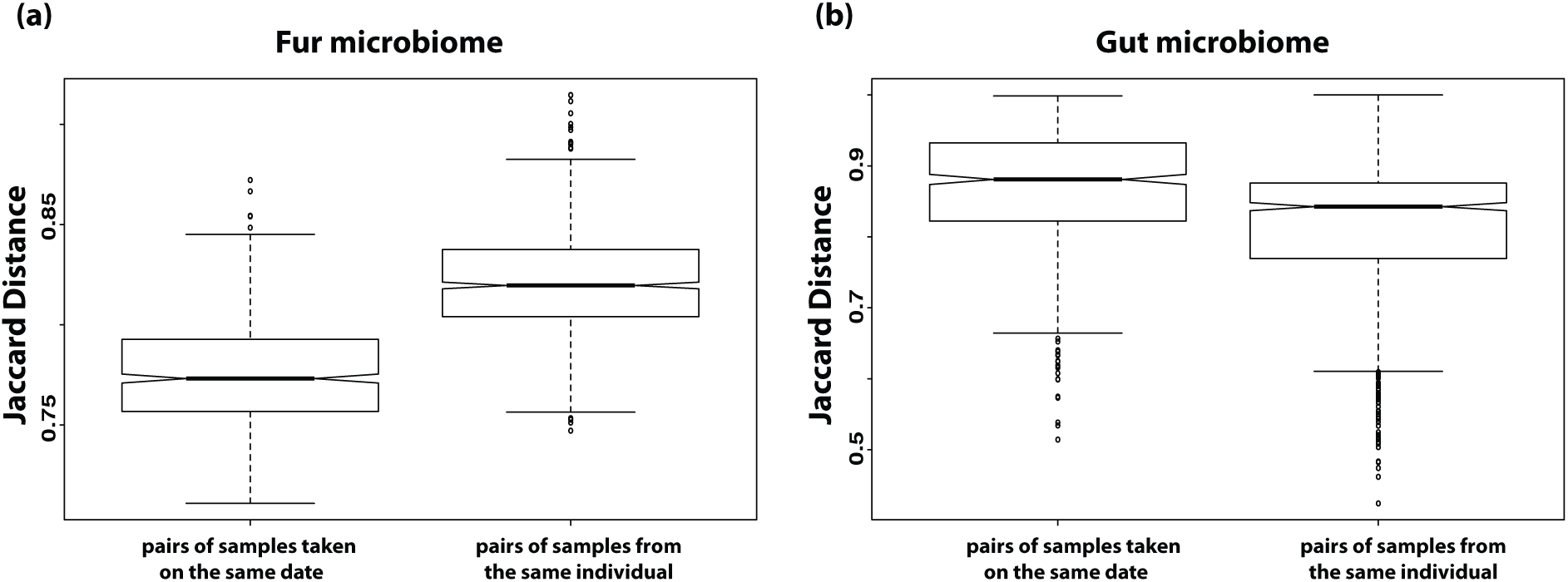
Similarity of pairs of samples in the experiment colony, from the same individual or from the same date. In the fur microbiome **(a)**, pairs of samples from different individuals, collected on the same date, are more similar than pairs of sample from the same individual from different dates. In the gut microbiome **(b)**, pairs of gut samples from different individuals, collected on the same date, are less similar than pairs of samples from the same individual on different dates. Box plots show the median and distribution of the Jaccard distance between all pairs of samples in each of these categories; hinges represent distribution quartiles, and notches the 95% confidence intervals of the medians. Differences between all distributions are highly significant (Kruskal-Wallis test, p<0.0001).

This finding is supported by a number of additional analyses (see Supplementary Sections 2 and 3): (1) Qualitatively similar results are obtained when using different weighted and unweighted distance measures (Binary, Bray-Curtis, Jaccard, Jensen-Shannon divergence, unweighted Unifrac, and Weighted Unifrac), and considering different bacterial taxonomic levels. (2) The pattern of coordinated change of the fur microbiome across the whole colony was even clearer when we included samples from the entire colony. The colony consisted of 33 individuals, each of which was sampled once at the beginning of the experiment and again at its end. (3) Results were robust to multiple conservative data filtering schemes, which ensured that possible bacterial contamination had been removed from the dataset (see Methods and Supplementary Section 2).

These colony-level changes over time are not easily explained by the study of the dynamics of particular microbial taxa (see examples of such dynamics in supplementary Section 4). Instead, the colony-level dynamics seem to be an emergent property of the host-microbiome system as a whole. The dynamics can be observed most clearly when considering the overall composition of the bats’ microbiomes. They are most obvious when the microbiome composition is measured in terms of only the presence or absence of each taxon and not their relative abundances, suggesting that a prominent part of the change in time occurs in microbial species that are generally found at low frequencies (see Supplementary Section 2).

Fur microbiome samples from the open colony, collected on the same dates, do not share the temporal trajectory of the experimental colony, ruling out the possibility that the interindividual similarities result from artifacts in the collection or sequencing processes (Supplementary Section 1). The samples from the open colony do, however, recapitulate the pattern found in the experiment colony: they show a significant colony-level change over time, supporting the generality of the finding and showing that it is not an artifact of captivity (Supplementary Section 1, Figures S1.1-S1.2; sample date explains 60% of the variance and individual identity explains 10% of the variance according to PERMANOVA; p<0.001). The fur microbiomes of the bats in the two colonies were similar: 96% of the species found in the open colony were also found in the experiment colony. However, microbial alpha diversity was somewhat higher in the experiment colony, with mean Shannon index of 6.39 in the experimental colony, and 5.97 in the open colony (a statistically significant difference: two-tailed t-test, p<0.001; calculated on presence/absence data, at the OTU level, on the dataset cleaned from potential contaminants; see additional measures and further details in supplementary Section 6).

A parallel analysis of the gut microbiomes yielded a different pattern: although sampling date was found to be a statistically significant explanatory variable, explaining circa 10% of the variance among samples (PERMANOVA, p<0.001), it was secondary to individual identity, which explained approximately 30% of the variance (PERMANOVA, p<0.001). Accordingly, in agreement with the findings reported for a range of body sites in humans and other vertebrates (24, 25, 29, 40), pairs of gut microbiome samples from the same individual are more similar to one another than pairs of samples from the same day but from different individuals (Figure 2b; Kruskal-Wallis rank sum test, p-value<0.001; confirmed using a Mantel test to avoid pseudo-replication, p-value<0.001).

The difference between the main factors driving the dynamics in the two body sites, date in the fur and individual in the gut, highlights the colony level dynamics as a feature not of the bat microbiomes in general, but of the bat fur microbiome specifically. This is true despite the fact that the diet of all individuals in the captive colony was almost identical, a factor that should have increased the similarity of individuals’ gut environments and therefore their microbiomes.

The fur microbiome is expected to be strongly influenced by the fur chemistry, and also to influence that chemistry. To examine the correlation between fur microbiome and fur volatiles, we collected fur samples from the experimental bats every two weeks and analyzed the composition of their volatile molecules by gas chromatography (GC). We found a pattern analogous to the one seen in the fur microbiome: the prominent factor governing variability is a change in the volatile profile over time, which is common across individuals (**Figure 4a**; Adonis PERMAONVA, variance explained: 27%, p<0.001). As in the case of the microbiome, individual identity is less important in explaining the composition of samples and it does not reach a significance threshold in a PERMANOVA test (p=0.43; see also **Figure 4b**).

**Figure 4:**
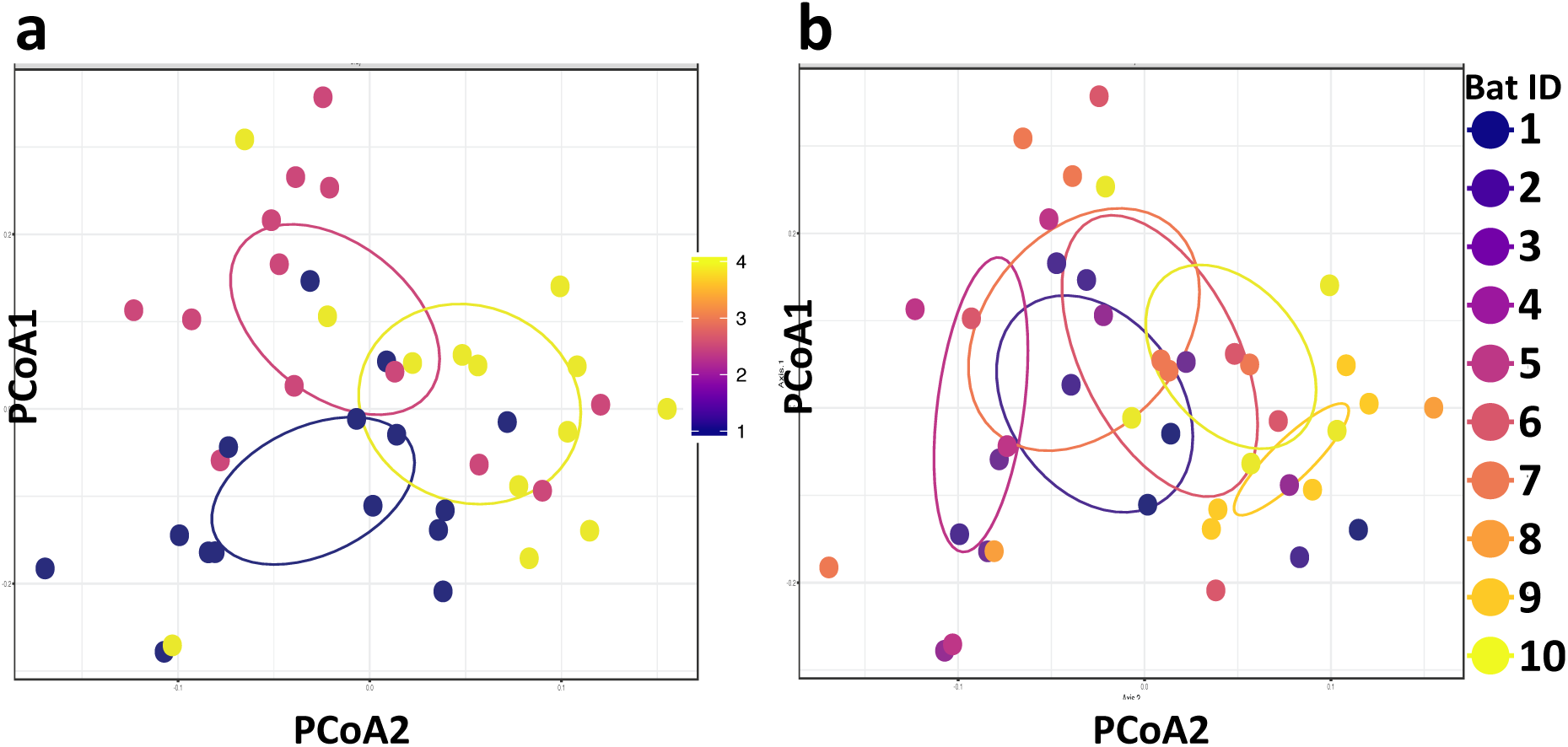
Colony and individual-level patterns in volatiles’ profiles. Samples of the profile of volatile compounds in the fur of the ten focal bats in the experiment colony, plotted using PCoA of the Bray-Curtis dissimilarity measure between samples. Each point represents a sample. **(a)** Samples are colored according to collection date; dates are divided into the three time trimesters of the 13-week period of the experiment (trimesters 1-3 are colored blue, red, and yellow, respectively). Only 3 periods (vs. 4 for the microbiome) were used for volatile analysis due to the smaller number of samples. **(b)** Samples are colored according to the individual from which they were collected. Ellipses represent the areas around centers of mass (see Methods).

The concentrations of a number of volatile compounds were significantly correlated with the abundance of certain bacterial taxa in the fur, which are known producers of these volatile metabolites: Cholestane diene was significantly positively correlated with three taxa of the order Actinomycetales (genera *Nesterkonika*, *Arthrobacter* and *Brevibacterium*). Palmitic acid was significantly correlated with the genus Neisseria. Oleic acid was significantly correlated with the genera *Alkanindiges* and *Neisseria.* All three compounds are known to play a role in communication among vertebrates (see supplementary section 5). This suggests that some of the change in volatiles over time might reflect the respective colony-level changes in the fur microbiome (the volatile dynamics remained significantly dependent on time even when the dataset is reduced to include only these bacteria-related volatiles: Adonis PERMANOVA, variance explained: 20%, p<0.005,). It is likely that the two modalities – fur chemistry and microbiome composition – interact with one another.

## Discussion

Unlike findings in other vertebrates, the microbiome of the fur of the Egyptian fruit-bat changes over time in a manner that is coordinated across the whole colony; this coordination is the prominent driver of variation in our data. Why are the temporal dynamics of the Egyptian fruit bat fur microbiome different from those of microbiomes in other studied mammals (e.g. (26, 29))? We propose that the frequent physical interactions between the bats in a colony (the bats perch in very tight clusters) have a homogenizing effect on their fur microbiomes, leading to dynamics in which the fur microbiomes of all individuals in the colony function together as a single ecosystem or super-organism. The changes over time may be driven by external factors, such as changes in the physiological state, in diet, or seasonal changes (although such changes were largely controlled in our experiment; see supplementary Section 8), but also by processes that are “internal” to the bacterial community such as neutral drift, local adaptation, and ecological succession.

Similar dynamics to those we find have been described in datasets from individuals along a developmental trajectory such as studies of human infant microbiomes (41–43), suggesting an ecological succession process, driven by physiological maturation of the host. The vast majority of individuals in our study were fully mature, so this cannot be the underlying driver of the pattern we see. However, a physiological change of that nature may account for some of the microbiome change over time in our experiment colony: for example, changes in the females’ reproductive state, which were correlated across most females and that became pregnant at about the same time, accounted for 4% of the microbiome variation (PERMANOVA, Adonis method in R, p<0.0001).

Our second main finding is that the gut microbiome is not characterized by such a prominent change in time as seen in the fur. Why are the dynamics of the fur and the gut microbiomes so qualitatively different? One possibility is that the difference is due to the relative role that common environmental factors play in each of these two modalities: the fur environment is strongly influenced by external factors, while the gut environment is strongly affected by the individual’s physiology and immune system, which buffer it from such environmental influences as diet, which is largely common to all individuals in our colony. This buffering can be seen as adding a “personalizing” effect, increasing the role of individual identity determining gut microbiome composition. Another possibility is that the different dynamics stem from differences in dynamics of bacterial transmission: the bats’ behavior, which includes frequent and extensive physical contact, has a homogenizing effect on the fur microbiomes, a process from which the gut microbiome is relatively shielded. From the bacterial perspective, one can think of gut bacteria as facing a greater transmission limitation than fur bacteria, creating a structured meta-population in which each individual’s gut constitutes an “island”, allowing both neutral and selectively-driven divergence between the microbiomes in different guts. These possibilities are not mutually exclusive.

The functioning of the colony’s fur microbiome as a single, highly connected, ecosystem might have important implications on the behavior and ecology of bats and other social species that roost in close proximity. Analysis of the volatiles found on the bats’ fur suggests that the fur microbiome may play a role in maintaining the social structure of the colony by facilitating olfactory-based recognition of colony members. Analysis of the volatiles in the two colonies (experimental and wild) at a single time point revealed that the two differ (see supplementary Section 5).

Bats’ associated microbes have recently received much attention from two specific perspectives: the first views bats as potential reservoirs of zoonotic pathogens that may infect humans (44–47). The second focuses on the pathogens of the bats themselves, particularly on dynamics of the white nose syndrome (48, 49) a serious emerging bat epidemic in bats (50, 51). The highly correlated dynamics of the colony members’ fur microbiomes suggests that in bats, and perhaps more broadly – in social species that roost in great proximity – the resilience to some types of disease may be largely a colony-level trait, and less a feature of individuals. This has obvious implications, potentially influencing plans for intervention that would mitigate the effects of the white nose syndrome or minimize the prevalence of specific zoonotic pathogens.

From a more theoretical evolutionary standpoint, our findings suggest that selective pressures on and through the fur microbiome, in species that are characterized by frequent physical contact between individuals, may act most prominently at the colony level, and not at the level of the individual, as is commonly assumed. This implies that it may be highly informative to supplement the study of host-microbiome dynamics with a meta-community framework that incorporates hierarchically structured transmission dynamics and in which colonies are the entities whose fate is studied.

## Methods

### Data collection

Two major colonies of bats are held in the Tel Aviv University zoological garden facility. The first, denoted *the experimental colony*, consisted of 33 bats at the time of this study. The second, denoted the *open colony*, consisted of 35 free ranging bats that can fly out and come back as they wish. From the experimental colony, the same 10 focal bats, 5 males and 5 females, were sampled once a week for their gut and fur microbiome during March to June 2016. Additionally, 4 focal bats from the open colony were sampled at 10 time points for comparison (not all were present in all 10 time points, as happens in a free ranging bat colony); mean number of samples from each bat is 7).

All bats were handled with single use clean gloves and swabbed for DNA before other measurements were taken, in order to limit contamination. The samples were taken by sterile culture swab applicators (BD CultureSwab™) moistened with Ringer’s Solution. Fur sampling was done by sweeping the swab, back and forth, 10 times over each of four different sites: shoulders, arm pits, stomach and muzzle. Sampling the gut microbiome was done by holding the bat and squeezing the anus to extract transparent discharge. This discharge was collected by sterile culture swab applicators moistened with Ringer’s Solution. *Rousettus aegyptiacus* has a relatively short intestine, not differentiated into small and large parts and with no observed cecum or appendix (38); the duration of the intestinal pass is approximately 40 minutes (38, 39). As the bats were after their day-fast and the intestine was free of content, we suggest that this discharge well represents the core gut microbiome without using invasive or lethal techniques (see supplementary Section 6 for a comparison of the microbiome in these samples and in those found in the bats’ feces). All bats were sampled in the same way and in the same order. Additional environmental samples were collected from the fresh food plates, capture nets, and air. After sampling, the swabs were sealed in a sterile plastic container provided, and immediately taken for DNA extraction.

### DNA extraction and pyrosequencing

Genomic DNA was extracted from swabs using the PowerSoil© DNA isolation Kit (MoBio Laboratories), as recommended by the manufacturer. Extracted DNA samples were stored at –20°C. PCR amplification of the 16S rRNA gene was carried out with universal prokaryotic primers containing 5-end common sequences (CS1-341F 5’-ACACTGACGACATGGTTCTACANNNNCCTACGGGAGGCAGCAG and CS2-806R 5’-TACGGTAGCAGAGACTTGGTCTGGACTACHVGGGTWTCTAAT). Twenty eight PCR cycles (95°C 15 sec., 53°C sec. 15, 72°C 15 sec.) were conducted using the PCR mastermix KAPA2G Fast™ (KAPABiosystems); successful amplification was verified by agarose gel electrophoresis. Sample-specific barcodes and Illumina adaptors were added in 8 additional PCR cycles, and paired-end deep sequencing of the PCR products was performed on an Illumina MiSeq platform at the Chicago Sequencing Center of the University of Illinois. Sequencing depth ranged from 1589 to 30000 sequences per sample; to ensure data evenness, data were rarefied to an equal depth of 1500 sequences per sample.

### Data analysis

Demultiplexed raw sequences were quality filtered (PHRED quality threshold <20) and merged using PEAR (52). Sequences shorter than 380bp (after merging and trimming) were discarded. Data were then analyzed using the Quantitative Insights Into Microbial Ecology (QIIME, version 1.9) package (53) in combination with VSEARCH (54). Sequences were de-replicated and ordered by size before OTU clustering at 99% threshold; to reduce spurious formation of OTUs, singleton sequences were not allowed to form new OTUs. Chimeric OTUs were detected and discarded using UCHIME (55) algorithm against the gold.fa database. Taxonomy was assigned using UCLUST (56) against the QIIME default database (greengenes 13.8).

Analysis downstream from QIIME was done in R and in Matlab. Primary R packages used were Phyloseq, Vegan and Caret. Statistical tests were conducted using their implementation in these packages, with the following settings: **PERMANOVA**: Adonis{vegan}, permutations = 10,000. **Mantel Test**: mantel{vegan} method = Pearson, permutations = 10,000. **Linear Discriminant Analysis (LDA)**: lda{caret}. In the PERMANOVA tests no strata were used, and the effect of each variable, e.g. date of sampling, individual identity, sex, and age, was assessed separately. Note that some of these variables are co-linear. This procedure does not control for pseudo-replication, and thus a Mantel test was conducted to support assertions regarding significance of variables wherever possible, and PCoA clustering was used for visual demonstrations. PCoA plots were made using ordinate{phyloseq}, with the default settings. The ellipses which describe each group’s center of mass are used for ease of visualization of the center of the distribution of the points in that group, and reflect the 25% confidence level around the center of a fit of these points to a multivariate normal distribution. Mantel tests were used to assess whether pairs of samples from the same individual and those from the same date are more or less similar to one another; this was done by performing a Mantel test on the matrix of Jaccard distances and the matrix obtained by assigning 1 to pairs of samples from the same date and 2 to pairs of samples from the same individual (see also (29)).

All results in the main text from Figure 2 onwards are for the dataset composed of the focal individuals only, following the most conservative procedure of omitting potential contaminant taxa. This included the removal of all microbial taxa that occurred in the negative controls or in more than one of the samples of the bats’ food at a frequency above 0.2%. The samples from the food were collected before it was introduced into the colony, and thus any microbial taxa in them were viewed as potential contaminants. This procedure may have omitted taxa that were not contaminants, and so the analyses were repeated with the full dataset as well, to confirm that they yield the same qualitative results. Wherever meaningful, analysis with the full range of samples is included in the supplementary material. PERMANOVA tests and LDA analysis were done using the matrix of relative abundances of microbial taxa, and PCoA plots in the main text present Jaccard distances based on presence/absence of microbial taxa. Analogous analyses with additional distance measures are presented in the supplementary material.

### Analysis of volatile compounds in fur using gas chromatography

Fur samples were placed in 3ml vials containing dichloromethane, for a minimum of 7 days. The samples were sieved, extracts were transferred to new insert vials while the fur was removed, dried and weighed for each sample. Two internal standards (udecanal and ergosterol) of known concentration (0.01 ng/μL) were added to each extract. Samples were first analyzed by combined gas chromatography/mass spectrometry (GC-MS;GC 7890A, MS 5975C; Agilent) using an HP-5MS capillary column, that was temperature programmed from 60°C to 300°C at 10°C/min. Compounds were identified by their mass fragmentation and retention times compared with synthetic standards when available. Compound quantification across samples was thereafter performed by gas chromatography with flame ionization detection (GC-FID) (CP 3800; Varian) using a DB-1 fused silica capillary column (30 m × 0.25 mm i.d.), temperature programmed as above, using peak integration. 22 peaks in the normalized chromatograms (Supplementary Section 5) were identified using GC-MS as biological compounds (rather than artificial contaminations). After removal of samples that failed to produce data, this process resulted in a matrix of 22 by 41 representing 22 volatiles sampled from 10 individuals over 6 time points (19 samples yielded no peaks probably because too little fur was collected and thus we had 41 and not 60 samples). Analysis of the resulting dataset was executed, for consistency, using the same methods and scripts as used for the PCoA and PERMANOVA analyses of the microbiome data. Correlations between abundance of microbial taxa and volatile compounds were carried out at the OTU level. For each of the 22 volatiles, a 41-dimentional vector representing the levels of this volatile across individuals and times was created. This vector was then (Pearson) correlated with a 41-dimentional vector representing the levels of an OTU (sampled over the same individuals and dates). Only the 30 OTUs that appeared in at least 50% of the samples of all individuals were used. This procedure was repeated over all 22 volatiles and 30 OTUs resulting in a (30x22) correlation matrix. Significant correlations were chosen following an FDR correction for multiple comparisons.

## Data Accessibility

All data supporting the results reported in this study will be made available upon publication.

## Acknowledgements

We thank Andrew Letten, Po-Ju Ke, Marion Donald, Natasha Dudek, Will Van Treuen, Liz Costello, Merav Maor, and Tovit Simon for technical help and insightful comments. O.K is supported by the John Templeton Fund and the Stanford Center for Computational, Evolutionary, and Human Genomics (CEHG). This research was partially supported by the European Research Council (ERC-GPSBAT).

## Footnotes

1 The bacterial community in each sample was characterized using multiplexed 16S rRNA gene amplicon sequencing. See Methods.

